# Increased replication-associated single-stranded DNA promotes formaldehyde-induced mutagenesis

**DOI:** 10.64898/2026.04.24.720578

**Authors:** Thomas Blouin, Charlotte McGuinness, Kierra Marshall, Caroline Bazzle, Natalie Saini

## Abstract

Formaldehyde (FA) is an environmentally abundant and endogenously produced aldehyde that has been shown to cause DNA damage, mutagenesis, and carcinogenesis. Several studies have demonstrated that FA induces guanine mutations resulting in a mutation signature like SBS40. In this work, we demonstrate that replication defects generating single-stranded DNA (ssDNA) caused by the downregulation of the major replicative polymerases results in elevated FA mutagenesis. We found that loss of Mrc1 (CLASPIN) resulted in a high accumulation of ssDNA and FA mutagenesis, and that these phenotypes were not dependent on Mrc1’s checkpoint activity. Loss of DNA-protein crosslink repair results in elevated FA sensitivity with no alteration to mutagenesis, likely due to the inability of the fork to bypass unprocessed protein adducts. Finally, we show that FA-induced mutagenesis is dependent on Pol ζ-mediated translesion synthesis, while deficiencies in the template switching pathway do not alter error-free bypass of FA adducts. Overall, our work points towards replication-associated ssDNA as a major substrate for FA-induced damage and elucidates the pathways that function to prevent FA mutagenesis at replication forks.

## Introduction

Formaldehyde (FA) is an environmentally abundant aldehyde and produced by cells endogenously through processes like 1-carbon metabolism and histone demethylation (Aravind and Koonin, 2001; Garrow et al., 1993; Kikuchi et al., 2008; Morellato et al., 2021; Porter et al., 1985; Shi et al., 2004; Thapa and Chan, 2025; Tsukada et al., 2006; Vijayraghavan and Saini, 2023a). Studies in rodents have shown that FA consumption either through inhalation or drinking water directly causes forestomach papilloma, leukemias, and gastrointestinal tract tumors (Kawanishi et al., 2014). Human population studies have also shown that occupational FA exposure is associated with increased mortality from myeloid leukemias and nasopharyngeal cancers (Beane Freeman et al., 2009; Coggon et al., 2003; Hauptmann et al., 2009, 2004; Pinkerton et al., 2004; Thapa and Chan, 2025). For these reasons, the International Agency for Research on Cancer has deemed FA a class I carcinogen (Kawanishi et al., 2014).

Cellular tolerance of FA falls broadly into two tiers, detoxification of the aldehyde and repair of its adducts (Dingler et al., 2020a). Essential to FA detoxification are the enzymes *ALDH2* and *ADH5,* and loss of these enzymes in mice has been shown to lead to disruptions in hematolymphoid development (Dingler et al., 2020b). Aberrant ALDH activity or expression has been found in breast cancer, colorectal cancer, lung cancer, head and neck squamous cell carcinoma, prostate cancer, pancreatic cancer, bladder cancer, and glioblastoma (Xia et al., 2023). High expression in breast cancer, head and neck squamous cell carcinoma, cervical cancer, and ovarian cancer has been associated with poor overall survival (Xia et al., 2023). It is likely that these cancers, due to higher metabolic requirements and proliferation, are required to upregulate these enzymes to tolerate larger endogenous aldehyde loads.

FA is a highly reactive electrophile that can form a variety of adducts on DNA including direct base adducts (mostly N^2^-HOMeG and N^6^-HOMeA), DNA interstrand crosslinks, and DNA intrastrand crosslinks (Beland et al., 1984; Cheng et al., 2008, 2003; Lu et al., 2012; Moeller et al., 2011; Zhong and Que Hee, 2005). FA is also known for its ability to form DNA-protein crosslinks (DPCs), most often with lysine and cysteine residues (Blouin and Saini, 2024; Lu et al., 2010). Cells resolve this damage through a variety of DNA repair pathways, including base excision repair, nucleotide excision repair, and homologous recombination (Kawanishi et al., 2014). Additionally, cells have evolved the ability to process DPCs directly at replication forks through the Wss1 (SPRTN) metalloprotease, further guarding against FA-induced genomic instability (Duxin et al., 2014a; Lopez-Mosqueda et al., 2016; Noireterre et al., 2023; Perry and Ghosal, 2022; Reinking et al., 2020a; Ruggiano and Ramadan, 2021; Stingele et al., 2014; Vaz et al., 2016; Yang et al., 2017). Interestingly, coupling of detoxification defects with defects in the Fanconi Anemia pathway that processes interstrand crosslinks results in the development of acute T-cell leukemia (Dingler et al., 2020b; Garaycoechea et al., 2012; Langevin et al., 2011; Pontel et al., 2015; Rosado et al., 2011).

ssDNA is generated throughout the cell cycle and dysregulated accumulation of ssDNA poses a serious risk to genomic stability in the presence of a genotoxic agent. Because ssDNA lacks a template strand required for most canonical repair, DNA adducts formed within ssDNA are often bypassed by the DNA damage tolerance pathways translesion synthesis (TLS) and template switching (TS) whereby cells can directly replicate past damage (Edmunds et al., 2008; Makridakis and Reichardt, 2012; Malik et al., 2020; Prakash et al., 2005; Sale, 2013). Accordingly, various studies have shown that UV (Burch et al., 2011), enzymatic deamination (Chan et al., 2015), alkylating damage (Hudson et al., 2023; Roberts et al., 2012a; Saini et al., 2020; Sakofsky et al., 2014; Yang et al., 2010), and oxidative damage (Degtyareva et al., 2019, 2013) can all target ssDNA and produce unique mutational patterns (Saini and Gordenin, 2020).

Prior work from our lab and others used an inducible subtelomeric ssDNA system in yeast to show that acute FA treatment caused an abundance of guanine mutations within ssDNA regions (Blouin et al., 2025; Thapa et al., 2022). FA-induced mutagenesis within ssDNA was shown to depend on TLS, and loss of TLS resulted in chromosomal arm loss events (Blouin et al., 2025). A major source of endogenous ssDNA in cells is DNA replication, when the CMG helicase unwinds the double helix at replication forks (Bochman et al., 2008; Coster et al., 2014; Douglas et al., 2018a, 2018b; Ilves et al., 2010; Lewis et al., 2022; Moyer et al., 2006; Noguchi et al., 2017; Seo and Kang, 2018; Sparks et al., 2019). Typically, generation of ssDNA at the fork is tightly regulated by the intra-S-phase checkpoint whose main signaling kinase Mec1 (ATR) becomes activated by the accumulation of ssDNA-bound RPA and works to suppress origin firing and slow replication progression (Doksani et al., 2009; Iyer and Rhind, 2017; Santocanale and Diffley, 1998; Zeman and Cimprich, 2014). Loss of this pathway or alterations to the normal replisome composition, like replicative polymerase downregulation or loss of critical replisome proteins (Mrc1/CLASPIN, Tof1/Timeless, Csm3/Tipin) have been shown to cause the formation of abundant replication-associated ssDNA that is prone to breakage and/or mutagenesis (Bhagwat et al., 2016; Grabarczyk, 2020; Hoopes et al., 2016; Lou et al., 2008; Nedelcheva et al., 2005b; Osborn and Elledge, 2003; Roberts et al., 2012a; Sui et al., 2020; Tourrière et al., 2005).

While our prior work established FA as a ssDNA mutagen and described how TLS and DPC repair coordinate to bypass FA-induced DNA damage, it is currently unknown which sources of endogenously produced ssDNA may be targeted for FA-induced mutagenesis. In this study, we investigated ssDNA generated at dysregulated DNA replication forks as a target for FA-induced mutagenesis, as well as the pathways protecting replication forks from FA-induced genomic instability. Overall, we find that fork dysregulation significantly increases FA-induced mutagenesis. Mrc1’s coupling function is required for preventing mutagenesis, while intra-S-phase checkpoint signaling is required for cellular survival following FA exposure. Finally, TLS but not TS contributes to the bypass of FA-induced adducts. Our work provides further insight as to which genomic substrates are at an elevated risk for aldehyde-induced genotoxicity.

## Results

### Downregulation of replicative polymerases increases formaldehyde-induced mutagenesis and sensitivity

*Saccharomyces cerevisiae,* like mammalian cells, utilize Pol ε, Pol δ, and Pol α to faithfully replicate the genome (Kawasaki and Sugino, 2001). To begin probing the effect of FA on replication-associated ssDNA, we first generated *S. cerevisiae* with dysfunctional replication by replacing the endogenous promoters of *POL2* (catalytic subunit of the leading strand Pol ε), *POL3* (catalytic subunit of lagging strand Pol δ), and *POL1* (catalytic subunit of Pol α) with tetracycline repressible *tetO_7_* promoters (Bellí et al., 1998). RT-qPCR of *POL2* and *POL1* showed expression was reduced to ∼50%, and *POL3* expression was found to be reduced to ∼30% 4-hours after doxycycline addition (Figure S1, Table S1) (Porcher et al., 2024). Use of the same *tetO_7_* promoter has been previously shown to result in a 10-fold reduction in protein levels of Pol3 (Saini et al., 2013).

Spot dilutions were used to assess the sensitivity of polymerase downregulation strains to FA. We did not observe strong growth defects of yeast strains with replicative polymerase downregulation, as is evident on the Yeast Extract, Peptone, and Dextrose (YPD) with doxycycline plate (Figure 1A). At 2 mM FA however, *TetR-POL2* strains appear slightly sensitive compared to the wildtype (Figure 1A). *TetR-POL3* and *TetR-POL1* display no apparent sensitivity to FA (Figure 1A).

**Figure 1:**
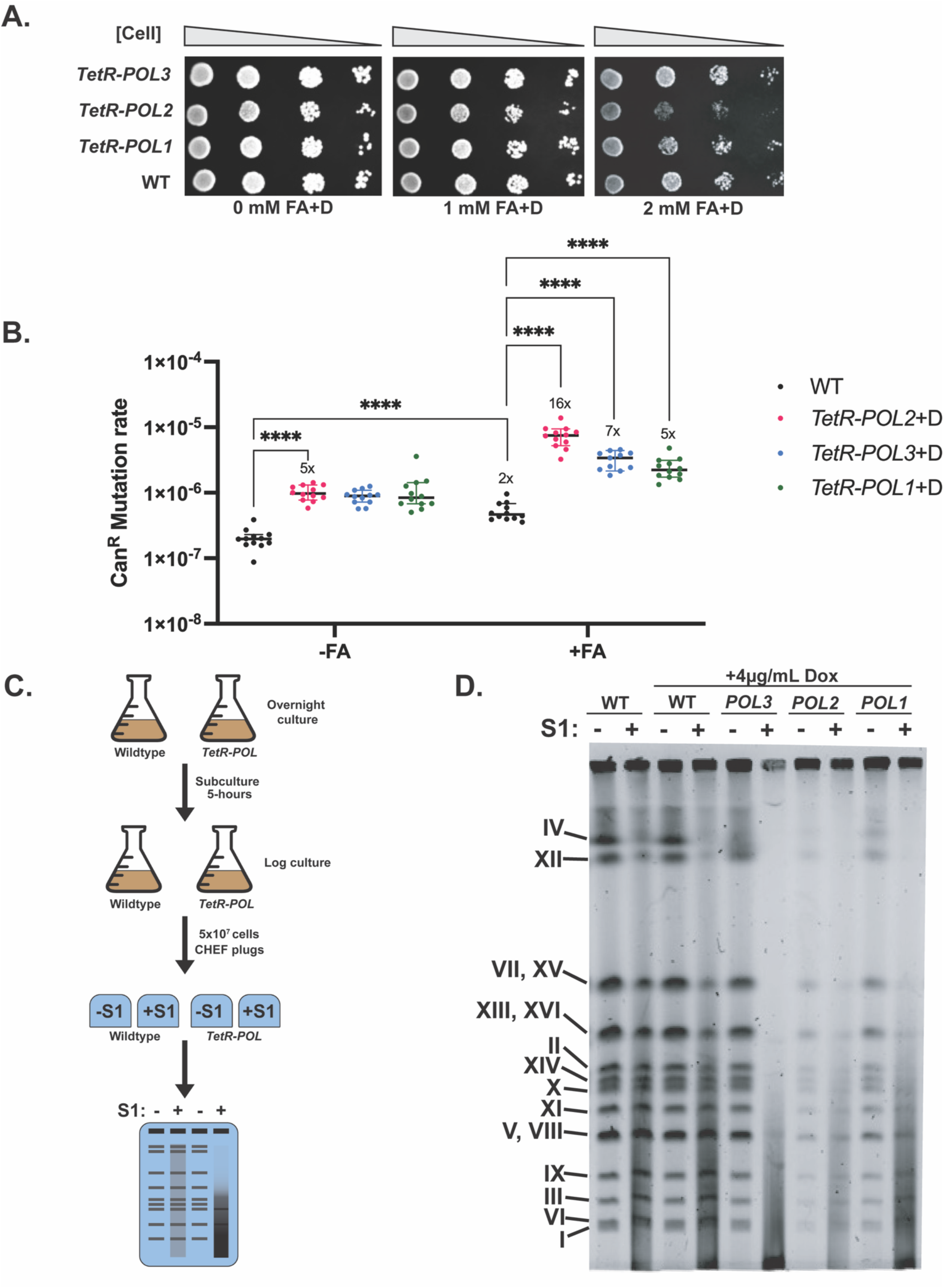
Downregulation of replicative polymerases increase FA-induced mutagenesis and sensitivity. A. Spot dilutions of replicative polymerase downregulation strains grown in the presence of 4 µg/mL doxycycline and various concentrations of FA for 2 days. B. Can^R^ mutation rate of various replicative downregulation strains when chronically exposed to 2 mM FA (+FA) or YPD (-FA) and 4 µg/mL doxycycline for four days. Bars represent median with 95% CI, and Y-axis is plotted on a log scale. P-values were calculated using a Mann-Whitney unpaired U-test and multiple testing correction was performed by analyzing stacked P-values. **** identifies q=0.0001. All significant changes with a greater than 2-fold difference are depicted on the graph. C. Experimental design for ssDNA detection using S1 nuclease and CHEF. All *TetR-POL* strains were grown in the presence of 4 µg/mL doxycycline. The representative CHEF gel shows expected results for a strain with low ssDNA in lane 2 and a strain with high ssDNA in lane 4. D. Representative CHEF gel of wildtype or *POL* downregulation strains. +/- corresponds to addition of the ssDNA specific S1 nuclease.

We further assayed FA-induced mutagenesis in each of these strains. The wildtype yeast strains demonstrated a significant but modest 2-fold increase in the Can^R^ mutation rate upon FA exposure (Figure 1B, Table S2). This suggests that in the absence of abnormal levels of ssDNA, FA is weakly mutagenic to *S. cerevisiae*. This assay was repeated using the polymerase downregulation strains. In the absence of FA, we observed significant 5, 5, and 4-fold increases in the *TetR-POL2, TetR-POL3* and *TetR-POL1* strains respectively over wildtype (Figure 1B, Table S2). This data supports several reports that depletion of major replicative polymerases and overall weakening of replication forks results in elevated genome instability (Lemoine et al., 2005; Mertz et al., 2017; Zheng et al., 2016). Upon treatment with 2 mM FA and doxycycline, *TetR-POL2* strains showed a 16-fold increase in the Can^R^ mutation rate over FA-treated treated wildtype strains, and an 8-fold increase over the untreated *TetR-POL2* strains (Figure 1B, Table S2). Similarly, *TetR-POL3* strains exposed to FA and doxycycline showed a 7-fold increase in mutation rate over FA-treated wildtype strains and a 4-fold increase over untreated *TetR-POL3* strains (Figure 1B, Table S2). Finally, FA and doxycycline-treated *TetR-POL1* strains showed a 5-fold increase above FA-treated wildtype isolates and a 3-fold increase above untreated *TetR-POL1* strains (Figure 1B, Table S2). Taken together, these results suggest that FA is highly mutagenic in strains with dysfunctional replication forks.

To confirm ssDNA was being generated in the polymerase downregulation strains, we analyzed genomic DNA via contour clamped homogenous electric field electrophoresis (CHEF), wherein intact chromosomes can be separated as distinct bands, while ssDNA appears as diffuse smears. We used log-phase cultures growing in the presence of doxycycline to make CHEF plugs (Figure 1C). Each plug was split and half was incubated with the ssDNA specific S1 nuclease for 1.5 hours. Cleavage of chromosomes at replication-associated ssDNA gaps is expected to appear as a smear in the gels. S1 treatment of wildtype strains grown in either YPD or YPD supplemented with doxycycline resulted in minimal smearing on the gel and overall conservation of band peaks in the intensity plots, showing that wildtype strains generate low levels of ssDNA (Figure 1D, Figure S2A-D). In contrast, each of the polymerase downregulation strains showed extensive smearing and loss of chromosomal banding pattern upon treatment with S1 nuclease (Figure 1D, Figure S2E-J). Interestingly, CHEF analysis of 2 mM FA-treated isolates did not demonstrate a detectible increase in ssDNA above what was observed in the untreated wildtype or *TetR-POL2* strains (Figure S3A and B). These observations indicate that FA is not likely the source of ssDNA in the replication defective strains and causes damage to pre-existing ssDNA.

While the *TetR-POL2* strains grown in YPD containing doxycycline maintained a banding pattern like wildtype, we noted that the chromosome bands were consistently fainter than wildtype (Figure 1D). This was despite using equal number of cells to prepare the plugs. We posit that this lack of signal may be due to extensive replication fork stalling and ssDNA making branched intermediates that do not efficiently migrate into the CHEF gel. We did not see this decrease in chromosome intensity in the *TetR-POL1* and *TetR-POL3* strains, likely indicating lower levels of ssDNA, replication stalling, and branched intermediates being formed in these strains as compared to *TetR-POL2* strains. Overall, these results show that downregulation of the replicative polymerases results in higher ssDNA levels than what is observed in wildtype strains.

### Mrc1 function in replisome coupling is required to prevent formaldehyde-induced mutagenesis

Mrc1 is an integral component of the replication fork protection complex with dual roles as both a signaling adapter protein in the intra-S-phase checkpoint and as a physical link between Pol ε and the CMG helicase (Alcasabas et al., 2001; Chen and Zhou, 2009; Gambus et al., 2006; Hodgson et al., 2007; Katou et al., 2003; Lou et al., 2008; Nedelcheva et al., 2005a; Osborn and Elledge, 2003; Pardo et al., 2017; Szyjka et al., 2005; Tourrière et al., 2005; Yeeles et al., 2017). To investigate the role of Mrc1 in FA-induced mutagenesis, we performed fluctuation assays in the presence or absence of 2 mM FA. Similarly to the polymerase downregulation strains, we observed a significant 5-fold increase in mutagenesis in the untreated *mrc1Δ* strains when compared to the untreated wildtype, suggesting that loss of Mrc1 results in weakened forks (Figure 2A, Table S2). After chronic exposure to 2 mM FA, *mrc1Δ* strains were found to have a significant 10-fold increase in the Can^R^ mutation rate when compared to FA-treated wildtype strains, and a 5-fold increase in mutation rate over the untreated *mrc1Δ* strain (Figure 2A, Table S2). To determine whether Mrc1’s coupling or signaling function was required to prevent FA-induced mutagenesis, we reintroduced a previously described *MRC1-AQ* mutant allele into the *mrc1Δ* background (Osborn and Elledge, 2003). Importantly, this mutant can localize to the replication fork and maintain coupling between Pol ε and the CMG helicase, but is unable to be activated by Mec1 (ATR) upon replication stress (Osborn and Elledge, 2003). We found that mutation rates were not elevated in neither the untreated Mrc1-AQ strains nor following chronic exposure to 2 mM FA (Figure 2A, Table S3). Rad9 (53BP1) is another DNA damage checkpoint adapter protein that is typically activated by Tel1 (ATM) in response to double-stranded breaks but can compensate for loss of Mrc1 signaling leading to downstream Rad53 activation (Bacal et al., 2018; Emili, 1998; Gilbert et al., 2001; Pfander and Diffley, 2011; Siede et al., 1993; Sweeney et al., 2005). To ensure the reduction in mutation rate seen in *MRC1-AQ* strains was not because of Rad9 compensation, we generated *rad9Δ* and *MRC1-AQ-rad9Δ* strains. We observed Can^R^ mutation rates like wildtype in *rad9Δ* isolates, and only a modest 2-fold increase in *MRC1-AQ-*m *rad9Δ* isolates exposed to FA (Figure 2A, Table S2). These results suggest that the ability of Mrc1 to maintain a coupled replisome is responsible for protecting against FA-induced mutagenesis.

**Figure 2:**
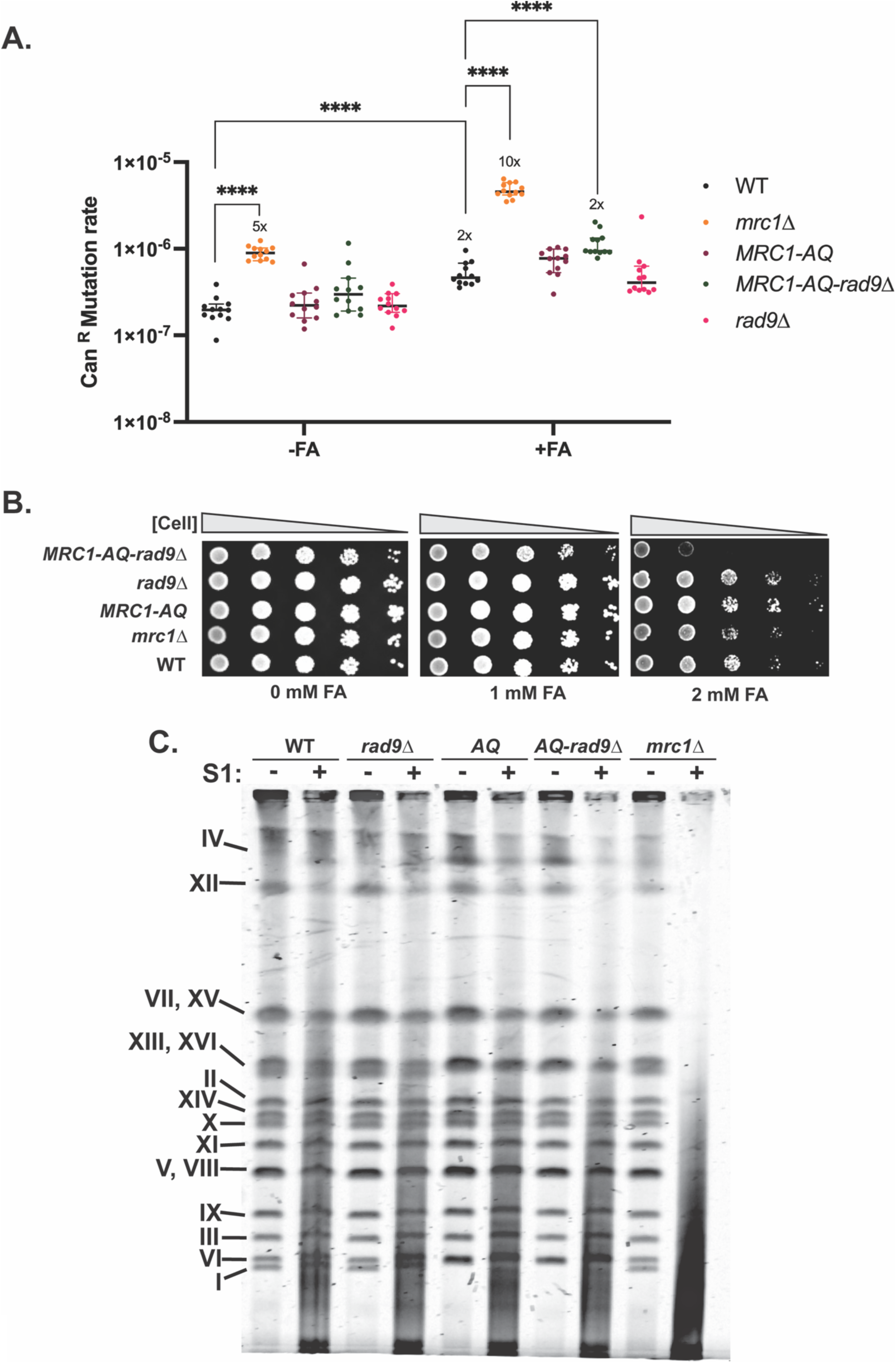
Mrc1 function in replisome coupling is required to prevent formaldehyde-induced mutagenesis. A. Can^R^ mutation rate of DNA damage checkpoint adapter protein-deficient strains when chronically exposed to 2 mM FA (+FA) or YPD (-FA) for four days. Bars represent median with 95% CI, and Y-axis is plotted on a log scale. P-values were calculated using a Mann-Whitney unpaired U-test. Multiple testing correction was performed by analyzing stacked P-values. **** identifies q=0.0001. All significant changes with a greater than 2-fold difference are depicted on the graph. B. Spot dilutions of checkpoint mutant strains grown in the presence of various concentrations of FA for 2 days. C. Representative CHEF gel of wildtype or checkpoint mutant strains. +/- corresponds to addition of the ssDNA specific S1 nuclease. *AQ* corresponds to *MRC1AQ* isolates.

Next, we asked if strains with ablated Mrc1 or Rad9 function were sensitive to FA. At 2 mM FA, *mrc1Δ* strains were observed to have a modest sensitivity relative to the wildtype (Figure 2B). *MRC1-AQ* and *rad9Δ* strains displayed no sensitivity, while the *MRC1-AQ-rad9Δ* strain displayed significantly heightened sensitivity. This result supports the idea that Mrc1 and Rad9 signaling works in parallel to maintain survival during FA exposure, likely by allowing efficient repair of FA-induced DNA adducts. Overall, our results suggest that Mrc1’s role in maintaining replisome coupling is essential in preventing FA-induced mutagenesis, and that DNA damage checkpoint signaling either via Mrc1 or Rad9 is required for cells to tolerate FA.

Finally, we assessed ssDNA levels in Mrc1 or Rad9 defective strains. We observed wildtype-like banding patterns in *rad9Δ, MRC1-AQ,* and *MRC1-AQ-rad9Δ* strains treated with S1, demonstrating that loss of checkpoint signaling adapters does not lead to accumulation of ssDNA (Figure 2C and Figure S4A-H). In contrast, we observed significant loss of the banding pattern and smoothing of the peak intensity profile in *mrc1Δ* strains, as expected given the role of Mrc1 in replisome coupling (Figure 2C and Figure S4I and J).

### DNA-protein crosslink repair promotes cellular survival to formaldehyde exposure but has minor impacts on mutagenesis

FA has been shown to be a potent DNA-protein crosslinker (DPC) (Blouin and Saini, 2024; de Graaf et al., 2009; Hoffman et al., 2015; Lai et al., 2016; Merk and Speit, 1998; Nakamura and Nakamura, 2020). In yeast, Wss1 (SPRTN) is a critical component of the cellular response to DPCs and has been shown to associate at DNA replication forks (de Graaf et al., 2009; Duxin et al., 2014a; Larsen et al., 2019; Lopez-Mosqueda et al., 2016; Mórocz et al., 2017; Noireterre et al., 2023; Reinking et al., 2020a, 2020b; Stingele et al., 2014; Vaz et al., 2016). To determine the impact of DPCs on FA-induced mutagenesis at dysregulated replication forks, we tested FA-induced mutagenesis in *wss1Δ, TetR-POL2-wss1Δ,* and *mrc1Δ-wss1Δ* strains using fluctuation assays. *wss1Δ* isolates showed no increase in mutagenesis above wildtype in untreated cultures or following chronic exposure to 2 mM FA (Figure 3A, Table S2). Spot dilutions revealed *wss1Δ* isolates to be significantly more sensitive to FA than wildtype isolates, underscoring the importance for Wss1 in cellular survival following FA exposure (Figure 3B). *TetR-POL2-wss1Δ* isolates displayed a minor 2-fold increase in Can^R^ mutation rate above *TetR-POL2* in the absence of FA, but no further increase was observed in the *TetR-POL2-wss1Δ* when grown in the presence of FA (Figure 3A, Table S2). Similarly, *mrc1Δ-wss1Δ* isolates had a comparable mutation rate to *mrc1Δ* both in the absence and in the presence of FA (Figure 3A, Table S2). Both *TetR-POL2-wss1Δ* and *mrc1Δ-wss1Δ* isolates appeared to be more sensitive to FA than the single mutants (Figure 3B). We conclude that an impaired ability to repair DPCs through loss of Wss1 has a minor impact on FA-induced mutagenesis both at wildtype and dysregulated replication forks. However, Wss1 is critical to cellular survival following FA exposure, resulting in a slight increase in sensitivity when DPC repair is lost in the context of dysregulated DNA synthesis.

**Figure 3:**
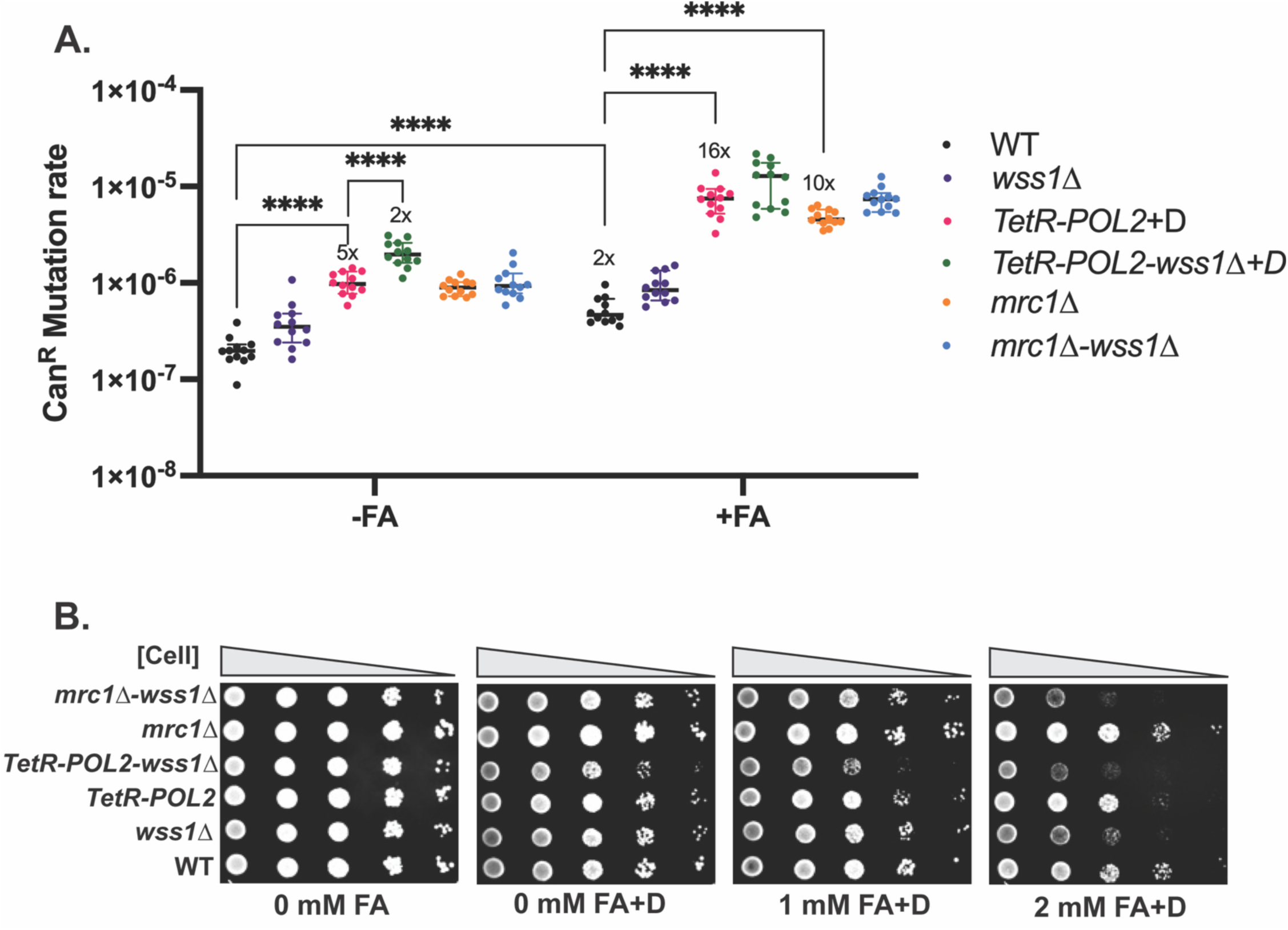
DNA-protein crosslink repair promotes survival following FA exposure but has minor impacts on mutagenesis. A. Can^R^ mutation rate of leading strand dysregulation and DPC repair defective strains when chronically exposed to 2 mM FA (+FA) or YPD (-FA) and 4 µg/mL doxycycline (*TetR-POL2* and *TetR-POL2-wss1Δ)* for four days. Bars represent median with 95% CI, and Y-axis is plotted on a log scale. P-values were calculated using a Mann-Whitney unpaired U-test and multiple testing correction was performed by analyzing stacked P-values. **** identifies q=0.0001. All significant changes with a greater than 2-fold difference are depicted on the graph. B. Spot dilutions of leading strand dysregulation and DPC repair defective strains grown in the presence of 4 µg/mL doxycycline and various concentrations of FA for 2 days.

### Template switching is not involved in the error-free bypass of formaldehyde adducts

Because ssDNA is not compatible with templated repair pathways, cells are increasingly dependent on DNA damage tolerance pathways when adducts evade repair and stall the replisome (Bainbridge et al., 2021; Frouin et al., 2003; Hedglin and Benkovic, 2017; Wong et al., 2021). Template switching (TS) uses homology on the complementary strand of an adjacent sister chromatid as the template for error-free DNA synthesis past an adduct (Ler and Carty, 2022). Alternatively, TS may involve the reversal of a stalled replication fork where nascent strands anneal with one another (Chang and Cimprich, 2009). TS depends on polyubiquitylation of PCNA which is carried out by Rad5 (HLTF), Mms2, and Ubc13 (UBE2), and deletion of any of these factors has been shown to equally impair polyubiquitylation (Hoege et al., 2002; Meister et al., 2025; Ulrich and Jentsch, 2000; Unk et al., 2006; Xu et al., 2016). To investigate the role of TS in bypassing FA-induced adducts, we generated *TetR-POL2-ubc13Δ, TetR-POL3-ubc13Δ, and ubc13Δ* strains. Loss of TS resulted in a significant 7-fold increase in the Can^R^ mutation rate in the absence of FA, suggesting that TS is critical to the error-free bypass of endogenous DNA adducts (Figure 4A, Table S2). Loss of TS in either fork dysregulation strain resulted in a further increase in the mutation rate of untreated cultures when compared to the fork dysregulation strain alone (Figure 4A, Table S2). Interestingly, deletion of *UBC13* both on its own and in the *TetR-POL2* and *TetR-POL3* backgrounds resulted in no significant increase in the Can^R^ mutation rate following FA exposure compared to untreated cultures (Figure 4A, Table S2). Spot dilutions revealed that loss of TS switching also did not increase sensitivity to FA (Figure 4B). Taken together, these results suggest that while TS is important in the error-free bypass of endogenously generated DNA damage, TS does not contribute to the bypass of FA-induced DNA adducts.

**Figure 4:**
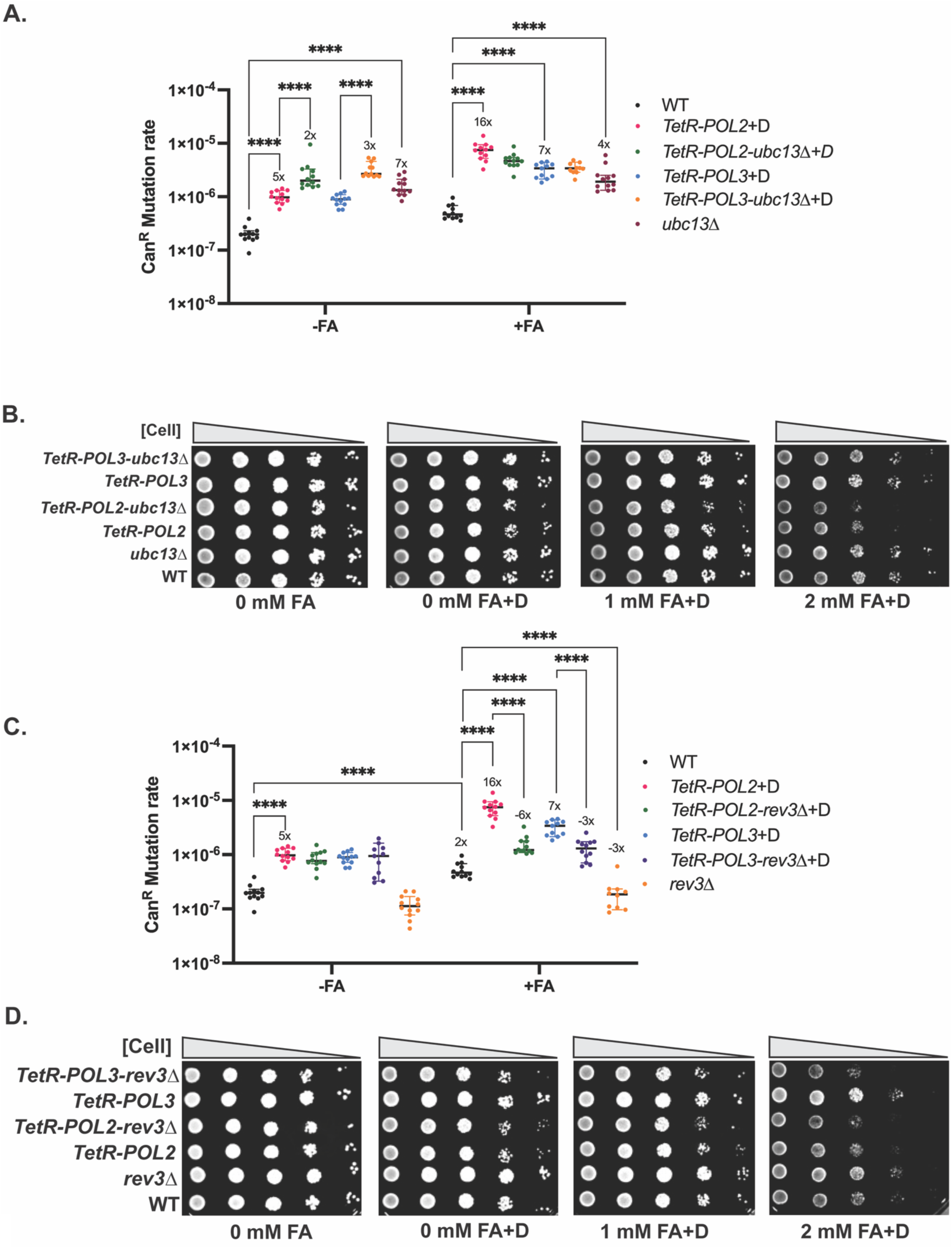
DNA damage tolerance pathways play differing roles in bypass of FA adducts. A. Can^R^ mutation rate of template switching deficient strains chronically exposed to 2 mM FA(+FA) or YPD (-FA) and 4 µg/mL doxycycline (*TetR-POL2, TetR-POL3, TetR-POL2-ubc13Δ,* and *TetR-POL3-ubc13Δ)* for four days. B. Spot dilutions of replication and template switching deficient strains grown in the presence of 4 µg/mL doxycycline and various concentrations of FA for 2 days. C. Can^R^ mutation rate of translesion synthesis deficient strains chronically exposed to 2 mM FA (+FA) or YPD (-FA) and 4 µg/mL doxycycline (*TetR-POL2, TetR-POL3, TetR-POL2-rev3Δ,* and *TetR-POL3-rev3Δ)* for four days. D. Spot dilutions of replication and translesion synthesis deficient strains grown in the presence of 4 µg/mL doxycycline and various concentrations of FA for 2 days. Bars in A and C represent median with 95% CI, P-values were calculated using a Mann-Whitney unpaired U-test and multiple testing correction was performed by analyzing stacked P-values. **** identifies q=0.0001. All significant changes with a greater than 2-fold difference are depicted on the graph.

### Translesion synthesis is required for the error-prone bypass of formaldehyde-induced DNA adducts

Translesion synthesis (TLS) function in yeast is dependent on Pol ζ (Haracska et al., 2003; Johnson et al., 2000; Martin and Wood, 2019; Prakash et al., 2005; Sale, 2013; Shachar et al., 2009). To investigate the role of TLS in bypassing FA-induced adducts, we generated strains lacking Rev3, which is the catalytic subunit of Pol ζ (*rev3Δ, TetR-POL2-rev3Δ,* and *TetR-POL3-rev3Δ*). Loss of TLS bypass did not alter the mutation rate of untreated strains, suggesting that TLS does not bypass endogenously formed DNA adducts. In contrast, loss of Rev3 led to an overall reduction in mutagenesis upon FA treatment; *TetR-POL2-rev3Δ* strains had a 6-fold reduction in the Can^R^ mutation rate when compared to *TetR-POL2*, while *TetR-POL3-rev3Δ* strains displayed a 3-fold reduction in the Can^R^ mutation rate when compared to *TetR-POL3* (Figure 4C, Table S2). Finally, we observed that FA-treated *rev3Δ* strains had a 3-fold reduction in the mutation rate when compared to wildtype, suggesting that TLS is responsible for most FA-induced mutations even in the absence of ssDNA accumulation (Figure 4C, Table S2). *rev3Δ* strains did not appear any more sensitive to FA than wildtype (Figure 4D). Interestingly, loss of TLS in the context of Pol2 or Pol3 depletion resulted in further sensitization to FA than what was observed in strains with intact TLS (Figure 4D). This result suggests that while cells are able to tolerate loss of TLS in the absence of abundant ssDNA, loss of TLS during fork dysregulation increases FA sensitivity likely due to increased break events as was shown previously (Blouin et al., 2025). Overall, our results suggest that FA-induced mutagenesis at dysregulated DNA replication forks depends on error-prone TLS bypass.

## Discussion

FA has been shown in various studies to induce mutations and genome instability. In *Saccharomyces cerevisiae*, a *lys2* frameshift reversion assay was used to show FA induces large, nucleotide-excision repair dependent deletions and established a role for TLS in the bypass of FA-induced adducts (Grogan and Jinks- Robertson, 2012; Vijayraghavan and Saini, 2023b). FA has been shown to mutagenize areas of ssDNA in yeast, resulting in an SBS40-like mutation signature that has also been observed in mice (Dingler et al., 2020b; Thapa et al., 2022). Recently, our lab confirmed these findings and established that FA-induced adducts were forming predominantly on guanine residues in ssDNA (Blouin et al., 2025). Pol ζ-mediated TLS was required for mutational accumulation within ssDNA and in the absence of TLS, there was a significant increase in the frequency of chromosomal arm loss events (Blouin et al., 2025). Efficient repair of these arm loss events was dependent on the DPC protease Wss1, suggesting that DPCs are a major adduct type within areas of ssDNA (Blouin et al., 2025). However, the above study utilized a *cdc13-1* mutant to generate ssDNA at sub-telomeric regions in yeast, resulting in a G2 arrest that allowed for an acute FA treatment. Further work is therefore needed to understand how these pathways may be involved in the FA-induced mutagenesis of endogenously generated ssDNA.

In this work, we demonstrate that replication-associated ssDNA is also highly susceptible to FA-induced mutagenesis. We find that dysregulation of DNA replication through the downregulation of the *POL1, POL2,* and *POL3* genes or deletion of *MRC1* causes an increase in ssDNA accumulation. Moreover, downregulated *POL1, POL2, POL3,* and *mrc1Δ* strains experienced elevated FA-induced mutagenesis. Such mutagenicity of ssDNA is likely due to the exposure and damage of primary amines typically protected within the interior of the helix, resulting in FA-induced lesion formation. These adducts typically persist due to the lack of template directed repair and must thus be bypassed to prevent prolonged polymerase stalling (Figure 5). Interestingly, we observed a stronger genotoxicity of FA in strains with *POL2* downregulation than with downregulated *POL3* or *POL1*. (Figure 1A and B). This increased mutagenic effect on the leading strand is potentially explained by the different repriming efficiencies of the leading and lagging strands.

**Figure 5:**
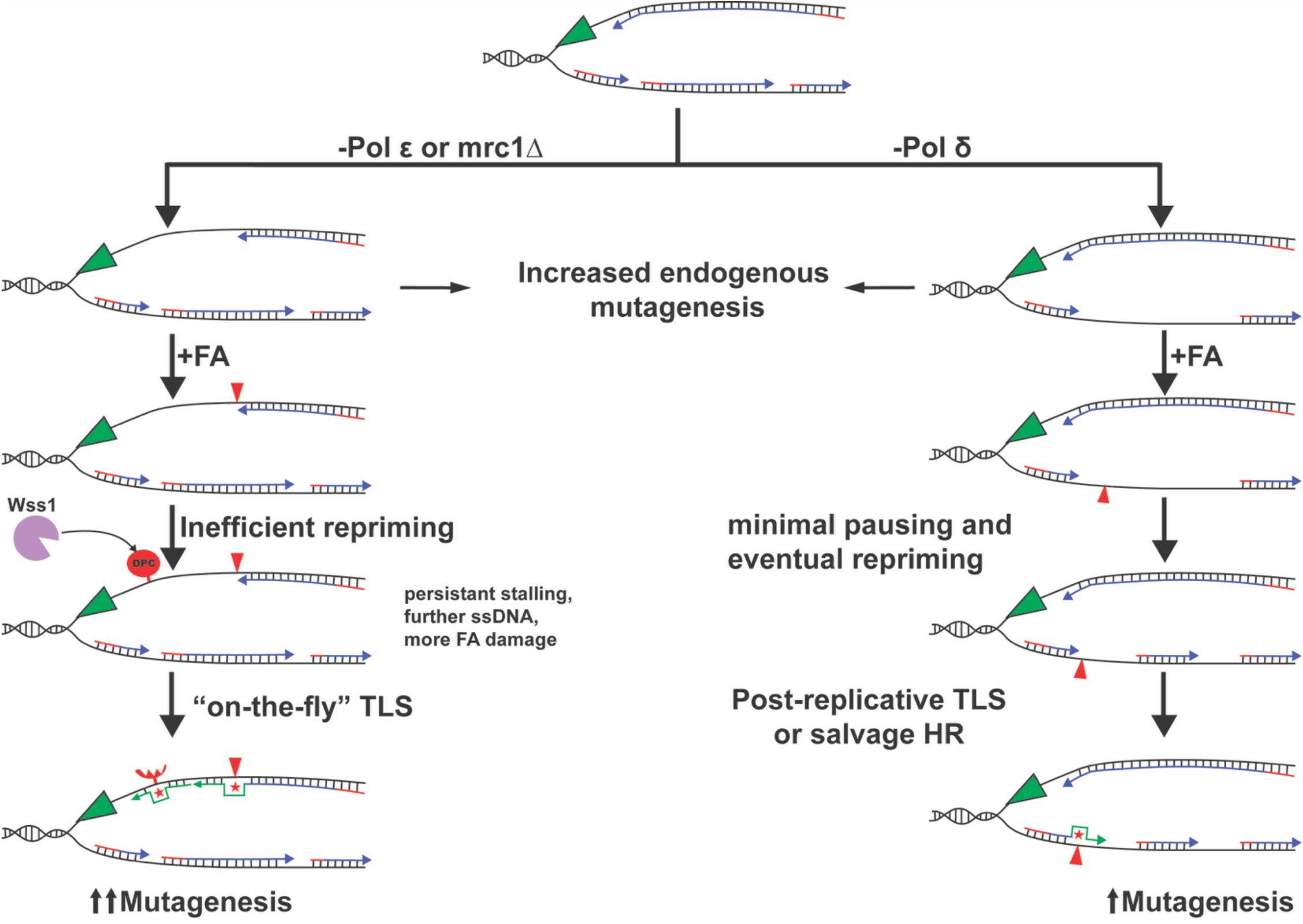
Proposed model for FA-induced mutagenesis at dysregulated leading and lagging strands. Downregulation of the replicative polymerases increases the amount of replication-associated ssDNA, and this dysregulation increases endogenous mutations. Leading strand adducts stall replication until TLS or repriming can occur. Because repriming of the leading strand is thought to be inefficient, leading strands exhibit a greater dependency on “on-the-fly” TLS or risk further uncoupling, more damage accumulation, and higher mutation rates. Adducts on the lagging strand also stall replication, but efficient repriming limits the generation of ssDNA at the fork, thus favoring a post-replicative model of TLS.

Eukaryotic leading strands are typically synthesized continuously from one RNA-DNA primer hybrid, limiting the need for repriming (Burgers and Kunkel, 2017). Upon collision with a Pol ε stalling lesion, the damage must either be bypassed by a DNA damage tolerance pathway or the leading strand may be reprimed downstream of the lesion leaving a daughter-strand gap to be dealt with in post-replicative repair (Bainbridge et al., 2021; Wong et al., 2021). While higher eukaryotes have acquired the primase PrimPol to more efficiently restart stalled leading strands, yeast lack this protein (García-Gómez et al., 2013). As such, reconstitution of the yeast replisome *in vitro* has revealed that leading strand repriming past an adduct is exceedingly rare and inefficient, likely placing a larger responsibility on “on-the-fly” DNA damage tolerance pathways to ensure stalling is temporary (Figure 5) (Taylor and Yeeles, 2019, 2018). Inefficient or delayed repriming of the leading strand has been shown to leave significantly larger ∼1600 nucleotide long gaps when compared to ∼200 nucleotide long gaps on the lagging strand (Hedglin and Benkovic, 2017; Lopes et al., 2006). Thus, potentially more FA-induced adducts could form on the leading strand resulting in the higher FA-induced mutation rate observed in *POL2* downregulation strains (Figure 1A, Figure 5). Further, lack of TLS would result in fork collapse and cell death (Figure 4D). Lagging strands however are synthesized discontinuously from many RNA-DNA primers, generating Okazaki fragments that must be ligated (Burgers and Kunkel, 2017). As such, repriming on the lagging strand happens quickly and limits the amount of uncoupling experienced (Taylor and Yeeles, 2018). This would result in smaller stretches of ssDNA and thus lower FA-induced mutagenicity and toxicity (Figure 1A and B). Depletion of Pol α has been shown to result in significantly longer Okazaki fragments on the lagging strand and an overall global decrease in replication origin firing efficiency (Porcella et al., 2020). This overall reduction in origin firing could limit the generation of ssDNA and potentially explains why Pol α strains had the lowest Can^R^ mutation rate in our assay (Figure 1A).

TS and TLS are critical DNA damage tolerance pathways that limit or overcome fork stalling (Bainbridge et al., 2021; Wong et al., 2021). In this study, we find that TS is critical for the error-free bypass of endogenous DNA damage but plays a minor role in bypassing FA adducts, while TLS plays a critical role in the error-prone bypass of FA-induced DNA damage (Figure 4). TS has been shown to bypass a variety of adducts, including UV damage, alkylation damage, and intrastrand crosslinks in an error-free manner (Fujiwara and Tatsumi, 1976; Jain and Siede, 2013; Masłowska and Pagès, 2022). Also, in yeast with extensive replication-associated ssDNA, TS was found to bypass abasic sites upon APOBEC3 or MMS-induced damage (Hoopes et al., 2016; Rosenbaum et al., 2019). Given that FA can induce a variety of similar adducts, the finding that loss of TS does not alter FA-induced mutation rates or sensitivity is surprising. It is possible that bulky FA-induced adducts prevent the strand exchange required for recombination-mediated TS. Further work is needed to determine the impact of template switching on FA-induced mutagenesis.

Loss of TLS was shown to significantly reduce the FA-induced mutation rate and further sensitize fork dysregulated strains to FA, suggesting that TLS is the major pathway responsible for the bypass of FA adducts (Figure 4C and D). These results support our prior findings that Pol ζ is required for the error-prone bypass of FA-induced adducts within areas of ssDNA, and that loss of this pathway can result in an increase in sensitivity, likely due to an abundance of double strand break events (Blouin et al., 2025). Furthermore, Pol ζ-mediated TLS is responsible for most FA-induced mutations even in the absence of fork dysregulation or abundant ssDNA (Figure 4C). This result might indicate that in replication proficient cells, even small regions of ssDNA are likely targets of FA-induced lesions and mutagenesis.

We observed that the presence of the checkpoint signaling adapters Mrc1 and Rad9 is crucial for cellular survival when exposed to FA, however, defects in this pathway did not alter mutagenesis by FA. We posit that FA likely generates mutagenic adducts on ssDNA, and fork stalling adducts ahead of the replication fork as well (Duxin et al., 2014b; Gao et al., 2023). The DNA replication checkpoint is essential to enable repair and bypass of such replication fork blocks. These blocks are not expected to cause increased mutagenesis and instead can culminate in double strand breaks and rearrangements if left unrepaired. Currently, our assay does not directly measure such events. A similar phenomenon was seen in strains deficient for DPC repair, pointing towards DPCs as a likely source of non-mutagenic fork blocking lesions (Blouin et al., 2025). Alternatively, adducts on ssDNA are bypassed by the error-prone TLS machinery resulting in Can^R^ mutagenesis. These data demonstrate that the DNA substrate alters the mutagenic outcome of FA exposure.

Because cancer cells are highly proliferative, they often have exposed replication-associated ssDNA. Various studies have reported DNA replication stress and accumulation of ssDNA gaps in cancer cell models, and these phenotypes are exacerbated in cells with replication or repair defects (da Costa et al., 2023a, 2023b; Falbo and Costanzo, 2026; Gaillard et al., 2015; Halazonetis et al., 2008; Karnitz and Zou, 2015; Wilhelm et al., 2020; Zeman and Cimprich, 2014). A key mechanism generating ssDNA gaps is the uncoupling of replicative helicases and polymerases (Bertolin et al., 2025; Falbo and Costanzo, 2026; Hashimoto et al., 2010; Kolinjivadi et al., 2017; Patel and Kim, 2023). Additionally, defective checkpoint activation can lead to repeated origin firing and drive the formation of ssDNA gaps even in the absence of fork stalling lesions (Bertolin et al., 2025). Cells lacking Pol ε subunits accumulate ssDNA gaps in a PrimPol and PARP1-inhibition dependent manner, likely due to the loss of the H3-H4 histone chaperone activities of Pol ε and PARP1 increasing fork stalling and repriming events (Falbo and Costanzo, 2026; Hill et al., 2024). Finally, loss of fork stabilizing proteins like BRCA2 or inhibition of ATR results in replication-associated ssDNA gap accumulation in human cells (Ortega et al., 2025; Situ et al., 2025). Such replication-associated ssDNA has been shown to be targeted by APOBEC3 enzymes, culminating in mutagenesis, genome instability, and cGAS-STING activation (Hoopes et al., 2016; Mertz et al., 2023; Ortega et al., 2025; Situ et al., 2025). Together, these studies highlight the importance of replication fork stability in preventing ssDNA accumulation, and the mutagenic potential of such ssDNA.

Several cancer types have been shown to have aberrant activity or expression of FA-detoxifying ALDH enzymes (Xia et al., 2023) It is likely that these cancers, due to higher metabolic requirements and proliferation, are required to upregulate these enzymes to tolerate larger endogenous aldehyde loads. Additionally, cancer cells, especially those deficient in *BRCA1/2* are known to carry large amounts of replication defects, ssDNA, and hypermutation (Chan et al., 2015; Cong et al., 2021; da Costa et al., 2024; Panzarino et al., 2021; Petljak et al., 2022; Simoneau et al., 2021; Taglialatela et al., 2021). These facts set up a potential therapeutic avenue whereby inhibition of ALDH enzymes increases endogenously produced aldehyde levels in cancer cells that damages the abundant ssDNA ultimately leading to cell death.

## Methods Yeast Strains

The strains used in this study were derived from ySR128 with the genotype *MATa, ura3Δ, can1Δ, ade2Δ, ura3Δ, his7-2, leu2-3, 112 trp1-289.* The genes *CAN1, URA3, ADE2* were reintroduced at the *LYS2* locus as the array *lys2::ADE2-URA3-CAN1* at a midchromosomal position of chromosome II (Roberts et al., 2012b). Genetically modified yeast used in this study were generated using homology-directed insertion of dominant drug resistance cassettes as previously described (Goldstein and McCusker, 1999). Strains were validated using PCR. The strains, primers, and plasmids used in this study can be found in Table S3. Strains carrying the *MRC1-AQ* allele are the same as described previously (Osborn and Elledge, 2003).

### RT-qPCR of polymerase downregulation strains

Cultures of wildtype and *TetR-POL1* yeast were grown in yeast extract, peptone, and dextrose (YPD) or YPD containing 4 µg/mL doxycycline (TCI America) overnight. The following day, these cultures were subcultured in fresh YPD and doxycycline for ∼4 hours and RNA was extracted using the Monarch Spin RNA Isolation Kit (New England BioLabs). RNA was quantified on a NanoDrop 1000 (Thermo Scientific) and quality assessed on a 1% agarose gel. Polymerase downregulation was measured via RT-qPCR using the New England Biolabs Luna Universal One-Step RT-qPCR kit. *ACT1* expression was used for normalization, generating Δ*Ct* values. Polymerase downregulation strains grown in YPD were compared to the wildtype strain grown in YPD, and polymerase downregulation strains grown in YPD with doxycycline were compared to wildtype in doxycycline, producing ΔΔ*Ct* values. Fold-change in gene expression was then calculated as 2*^−ΔΔCt^*. Gene expression levels of wildtype cultures were denoted as a fold-change of 1, and polymerase downregulation strains were compared against them. Primers used for RT-qPCR are described in Table S3b. RT-qPCR data for *TetR-POL2* and *TetR-POL3* were from Porcher et al., 2024, and primers used for these reaction can be found therein (Porcher et al., 2024).

### Fluctuation assay for measuring FA-induced mutagenesis

Strains carrying the *lys2::ADE2-URA3-CAN1* reporter array were streaked-out onto YPD (+4 µg/mL doxycycline for *POL* downregulation) plates or YPD plates containing 2 mM FA (Electron Microscopy Sciences) (+4 µg/mL doxycycline for Pol downregulation) and were incubated at 30°C for 72 hours. Individual colonies from the streak outs were patched onto fresh YPD plates with and without 2 mM FA and incubated for 24 hours. Two independent wildtype isolates were used to generate 6 patches, while two independent experimental isolates were used to generate 12 patches. Patches were resuspended in 1 mL of sterile water and used to create a dilution series. The 5^th^ and 6^th^ dilutions were plated onto synthetic complete (SC) media to determine the number of cells present in each patch, and the 0^th^ dilution was plated onto SC-Arginine plates containing 60 mg/mL canavanine (Sigma) and 20 mg/mL adenine to isolate Can^R^ colonies. Plates were incubated at 30°C for 72 hours and colonies were counted using an aCOLyte3 (Synbiosis Inc.). Mutation rates were calculated as previously described (Drake, 1991). Mutation rate figures show the mutation rate of each patch as a data point. The median is depicted as a horizontal bar and error bars depict 95% confidence interval. P-values were calculated using a Mann-Whitney unpaired U-test and multiple testing correction was performed by analyzing stacked P-values.

### Spot dilutions

Yeast strains were resuspended in water to a concentration of 1×10^7^ cells/mL. 200 µL of this solution was placed into the top wells of a 96-well plate. This solution was then diluted 10-fold per row of the spot dilution. 3 µL of solution from each well was spotted onto YPD, YPD with 4 µg/mL doxycycline, YPD with 1 or 2 mM FA, or YPD with both doxycycline and FA. Spots were grown at 30°C for 48 hours and imaged on aCOLyte3.

### Clamped Homogenous Electric Field Electrophoresis (CHEF)

CHEF gels were produced as previously described (Wei, 2006) in chapter 8 (PMID: 16118425). Strains were grown overnight in YPD or YPD containing 4 µg/mL doxycycline. The following day, cells were subcultured into a fresh 50mL of YPD with or without doxycycline and with or without 2 mM FA for ∼5 hours. Approximately 5×10^7^ cells were taken from each culture to create the agarose plugs. S1 digestion was performed the day the CHEF was ran by cutting one plug in half. Both halves were washed with S1 buffer (0.2 M NaCl, 50 mM sodium acetate, 1mM M zinc sulfate, 0.5% glycerol, pH 4.5) for 30 minutes 3 times. To one of the plugs, 500 u/mL S1 nuclease (Promega) was added, and both plugs were incubated at 37°C for 1.5 hours. For pulsed-field gel electrophoresis, a 1% agarose gel in 0.5X TBE was used and ran on a BioRad CHEF Mapper at a temperature of 14°C with switch time 60-120s, run time 24 hours, angle 120°, and voltage gradient of 6 V/cm. Gels were stained with ethidium bromide and visualized on a BioRad GelDoc Go imaging system. ImageJ (Version 1.54p) was used to generate the CHEF intensity plots. Briefly, this was done by inverting the background of the CHEF gel and using the rectangle tool to highlight lane 1 of the gel. The dimensions of this box were used to analyze all subsequent lanes. Once a lane was defined, the plot lanes feature was used, then analyze line graphs to get a .csv file of intensity values along the lane. The final intensity plots were generated in R.

## Author contributions

T.B, N.S conceptualized the study. T.B, N.S designed the experiments. T.B, C.M, K.M, C.B performed the experiments. T.B, C.M, K.M, C.B acquired the data, T.B and N.S analyzed the data. T.B and N.S wrote the manuscript. All authors reviewed the manuscript.

## Acknowledgements

The authors would like to thank Elizabeth Ampolini and Sriram Vijayraghavan for their critical reading of the manuscript.

---

**Figure S1:**
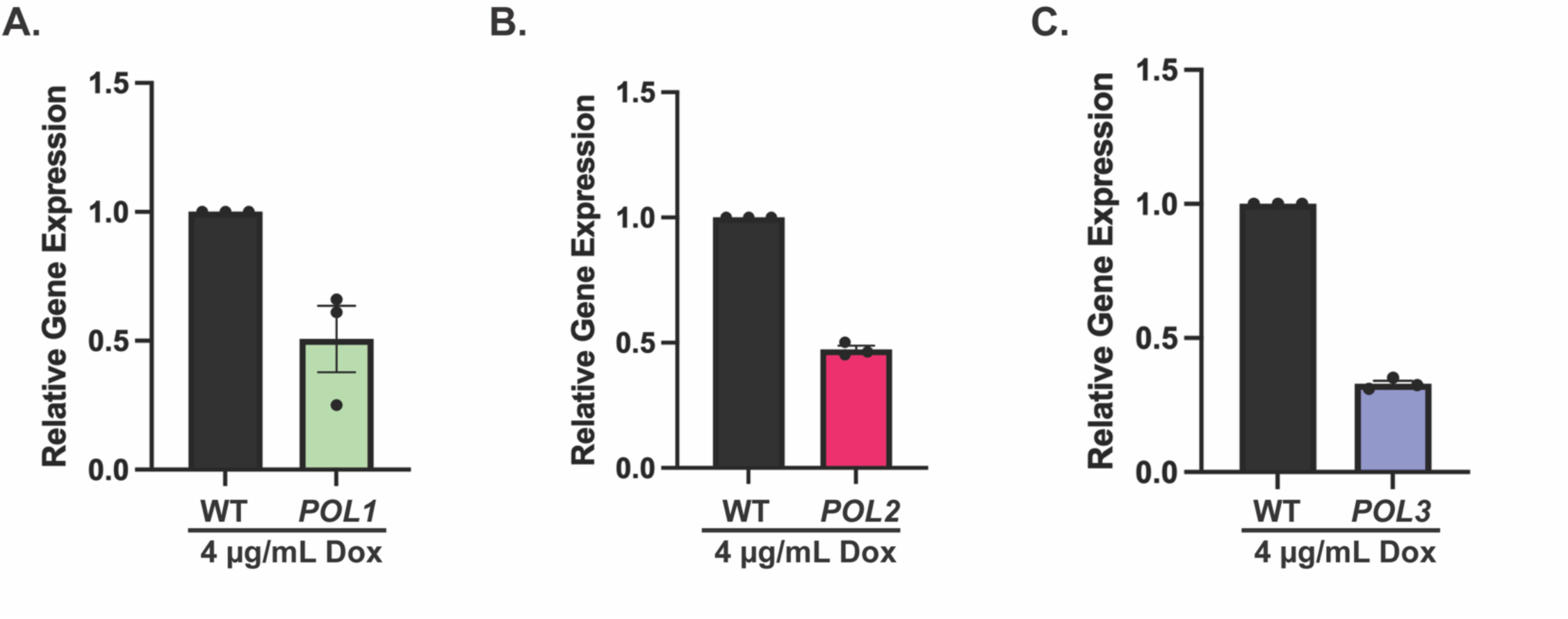
RT-qPCR of *TetR-POL* strains. Gene expression was monitored by RT-qPCR for the replicative polymerases under control of tetracycline repressible *tetO_7_* promoters. Expression was compared against the *POL* gene under the control of its endogenous (WT) promoter. Mean expression levels and standard error of the mean are depicted. Data for panels B and C are originally from Porcher et al., 2024.

**Figure S2:**
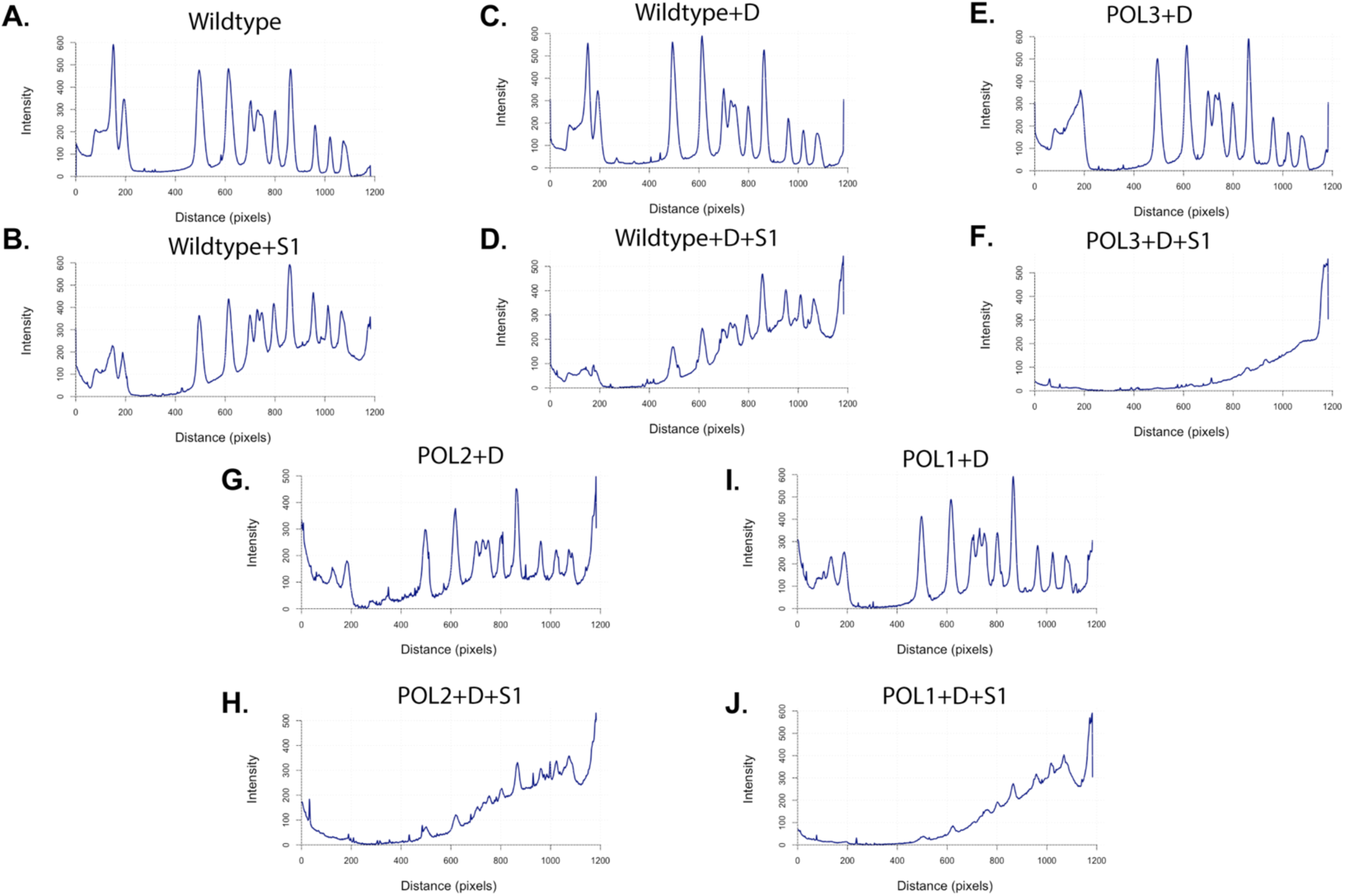
Peak intensity profile of polymerase downregulation CHEF gel. Peak intensity profiles of each lane in the CHEF gel featured in Figure 1D. Intensity plots were generated in ImageJ.

**Figure S3:**
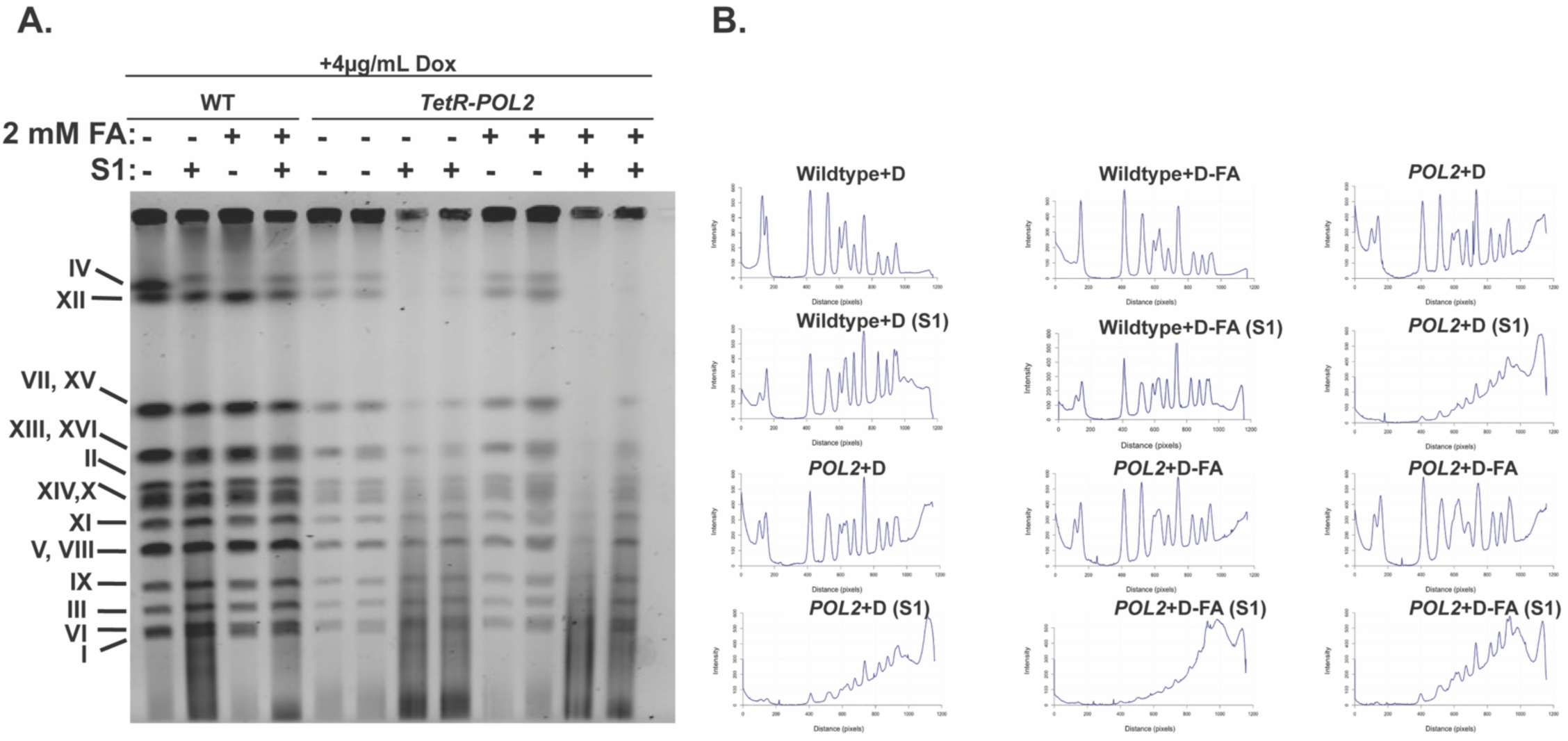
FA treatment itself does not result in higher levels of replication associated ssDNA. A. Representative CHEF gel of wildtype or *TetR-POL2* strains grown in the presence of doxycycline and/or FA exposed to the ssDNA specific S1 nuclease. B. Peak intensity profiles of each lane in the CHEF gel.

**Figure S4:**
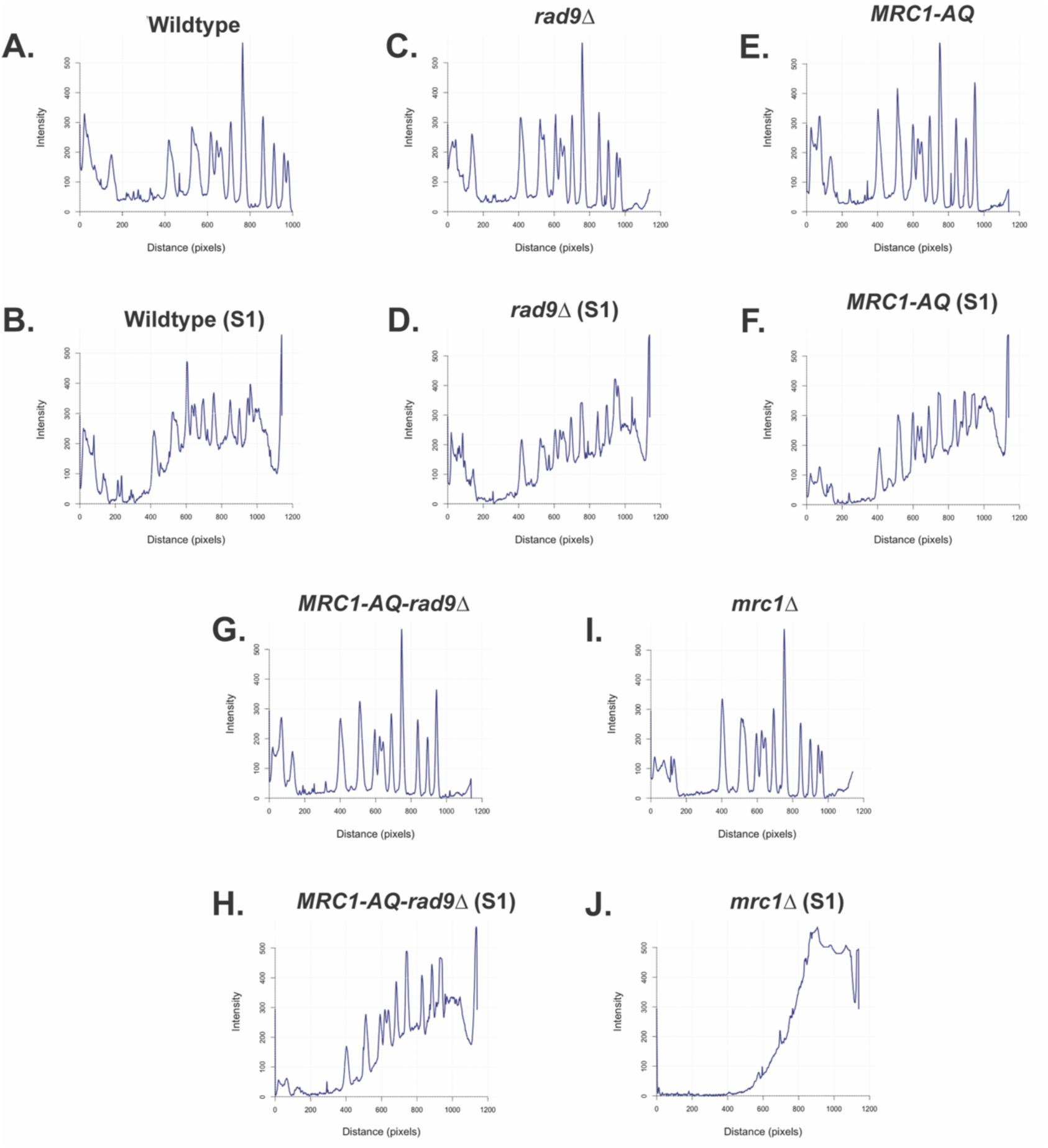
Peak intensity profile of Mrc1 and checkpoint signaling mutant CHEF gel. Peak intensity profiles of each lane in the CHEF gel featured in Figure 2C. Intensity plots were generated in ImageJ.

